# An Rshiny app for modelling environmental DNA data: accounting for false positive and false negative observation error

**DOI:** 10.1101/2020.12.09.417600

**Authors:** Alex Diana, Eleni Matechou, Jim E. Griffin, Andrew S. Buxton, Richard A. Griffiths

**Affiliations:** School of Mathematics, Statistics and Actuarial Science, University of Kent, Canterbury, UK; Department of Statistical Science, University College London, London, UK; Durrell Institute of Conservation and Ecology, School of Anthropology and Conservation, University of Kent, Canterbury, UK

**Keywords:** Bayesian variable selection, multi-level occupancy model, PCR, environmental DNA

## Abstract

1. Environmental DNA (eDNA) surveys have become a popular tool for assessing the distribution of species. However, it is known that false positive and false negative observation error can occur at both stages of eDNA surveys, namely the field sampling stage and laboratory analysis stage.
2. We present an RShiny app that implements the Griffin et al. (2019) statistical method, which accounts for false positive and false negative errors in both stages of eDNA surveys. Following Griffin et al. (2019), we employ a Bayesian approach and perform efficient Bayesian variable selection to identify important predictors for the probability of species presence as well as the probabilities of observation error at either stage.
3. We demonstrate the RShiny app using a data set on great crested newts collected by Natural England in 2018 and we identify water quality, pond area, fish presence, macrophyte cover, frequency of drying as important predictors for species presence at a site.
4. The state-of-the-art statistical method that we have implemented is the only one that has specifically been developed for the purposes of modelling false negatives and false positives in eDNA data. Our RShiny app is user-friendly, requires no prior knowledge of R and fits the models very efficiently. Therefore, it should be part of the tool-kit of any researcher or practitioner who is collecting or analysing eDNA data.

## 1 Introduction

Environmental DNA (eDNA) is increasingly used within biodiversity assessments (McClenaghan et al., 2020). The method relies on the detection of DNA released from source organisms into aquatic or terrestrial environments. This DNA is extracted from a sample of the substrate, usually water or soil (Thomsen and Willerslev, 2015) (stage 1), and then analysed using qPCR (Thomsen et al., 2012), or metabarcoding (Valentini et al., 2016) (stage 2).

We present an RShiny app for modelling single-species eDNA data by implementing the Bayesian model developed by Griffin et al. (2019). The model estimates site-specific probabilities of species presence while accounting for false positive and false negative observation error at both stages of eDNA surveys. The Griffin et al. (2019) model is an extension of the work by Guillera-Arroita et al. (2017) but in contrast to the latter, the Griffin et al. (2019) model does not require augmenting the eDNA data with other types of survey data. This is due to the specification of novel informative prior distributions that reflect our belief that the probability of a false positive observation is smaller than the probability of a true positive observation at either stage. Nevertheless, if opportunistic records of species presence exist for any of the sites, these can easily be accounted for within the model. Finally, Griffin et al. (2019) presented an MCMC algorithm that employs the Pólya-Gamma sampling scheme (Polson et al., 2013) and hence enables fast computation times and efficient Bayesian variable selection for all model parameters.

We give an overview of the Griffin et al. (2019) method in section 2 and present our RShiny app in section 3. A case study on great crested newt data collected by Natural England is presented in section 4 and the paper concludes with a discussion in section 5.

## 2 Materials and Methods

Griffin et al. (2019) presented a hierarchical Bayesian model that describes the different stages of eDNA surveys in terms of the probabilities of species presence and the probabilities of observation error. All of these probabilities can be functions of site-specific covariates, with dependencies modelled using logistic regressions. Subscript *s* is used to denote sites, *s* = 1,…, *S*, while subscript *m* is used to denote samples from sites, *m* = 1,…, *M*. The list of parameters is given in Table 1 and a schematic representation of the model is provided in Figure 1.

**Table 1:** Parameters of the Griffin et al. (2019) model. Note that *θ_01s_* = 1 - *θ_11s_*, *θ_00s_* = 1 - *θ_10s_*, *P_01s_* = 1 - *p_11s_*, *p_00s_* = 1 — *P_10s_*, and hence our RShiny app only reports results in terms of the probabilities of a positive (either true or false) observation at either stage.

| Name (probability of) | Detailed explanation (probability that) |
| --- | --- |
| $\psi_s$ species presence | site $s$ is occupied by the target species |
| Stage 1 |  |
| $\theta_{11s}$ stage 1 true positive observation | a sample from occupied site $s$ includes DNA of the target species |
| $\theta_{10s}$ stage 1 false positive observation | a sample from unoccupied site $s$ includes DNA of the target species |
| Stage 2 |  |
| $p_{11s}$ stage 2 true positive observation | a qPCR on a sample (from site $s$) that includes DNA of the target species is positive |
| $p_{10s}$ stage 2 false positive observation | a qPCR on a sample (from site $s$) that does not include DNA of the target species is positive |

**Figure 1:**
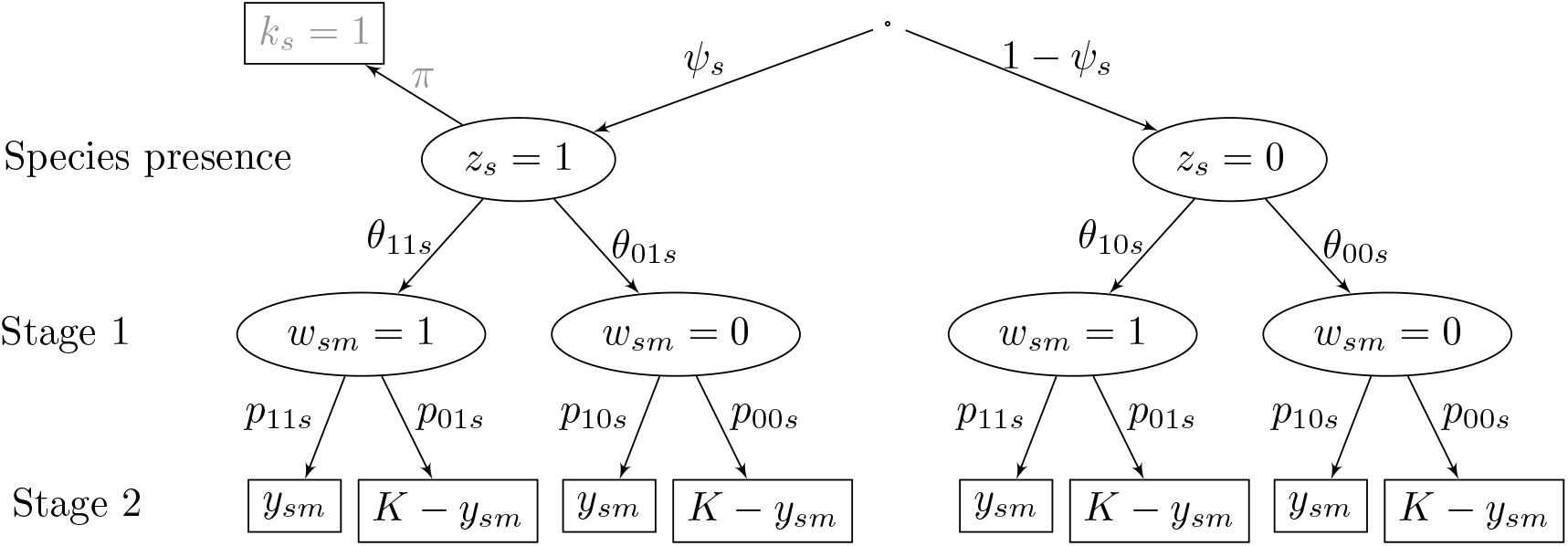
Schematic representation of the Griffin et al. (2019) model. Unobservable states are represented by ellipses and data by rectangles. The parameters are defined in Table 1. The latent variable *z_s_* indicates whether site s is occupied by the target species (1) or not (0) and the latent variable *w_sm_* indicates whether sample *m* from site s includes DNA of the target species (1) or not (0). The part of the model that is presented in grey corresponds to how opportunistic records of species presence are modelled. Specifically, parameter *π* indicates the probability that an occupied site has an opportunistic record associated with it and indicator variable *k_s_* indicates whether site s is known to be occupied (1) or not (0).

As explained in Griffin et al. (2019), the model is only locally identifiable, and this identifiability issue is overcome by introducing informative prior distributions that express our belief that a false positive observation is less likely than a corresponding true positive at each stage, so that the probabilities that *θ*_11_ < *θ*_10_ or *p*_11_ < *p*_10_ are small.

The MCMC algorithm presented in Griffin et al. (2019) employs the Pólya-Gamma sampling scheme (Polson et al., 2013), enabling fast computation for logistic regression models and efficient Bayesian variable selection, which is performed using an Add-Delete-Swap algorithm (Brown et al., 1998; Chipman et al., 2001). As the name of the algorithm suggests, at each MCMC iteration, we either propose to add a covariate that is not currently in the model, or to delete a covariate that is in the model, or to swap a covariate that is in the model with one that is not in the model at that iteration. This process gives rise to the posterior inclusion probabilities (PIPs), which indicate the proportion of iterations that each covariate was in the model for each parameter. PIPs can be used to understand how useful each of the covariates is as a predictor for the corresponding parameter and often a threshold of 0.5 is applied to identify the most important predictors (Ghosh, 2015).

## 3 RShiny app

The RShiny app is freely available and can be accessed via the RShiny server https://seak.shinyapps.io/eDNA/. However, we recommend that the app is downloaded via our dedicated website https://blogs.kent.ac.uk/edna/ and run locally.

The app includes a detailed help section that provides a step-by-step description on how to format the data, which need to be uploaded in a .csv file. Once the data have been uploaded, the user needs to specify the number of qPCR runs for each sample (this is assumed to be the same for all samples) and to select the parameters that are to be considered as functions of covariates, as shown in Figure 2. It is not necessary to consider covariates for any of the parameters, but the set of covariates uploaded will be considered as potential predictors for all parameters that have been specified in the settings window.

**Figure 2:**
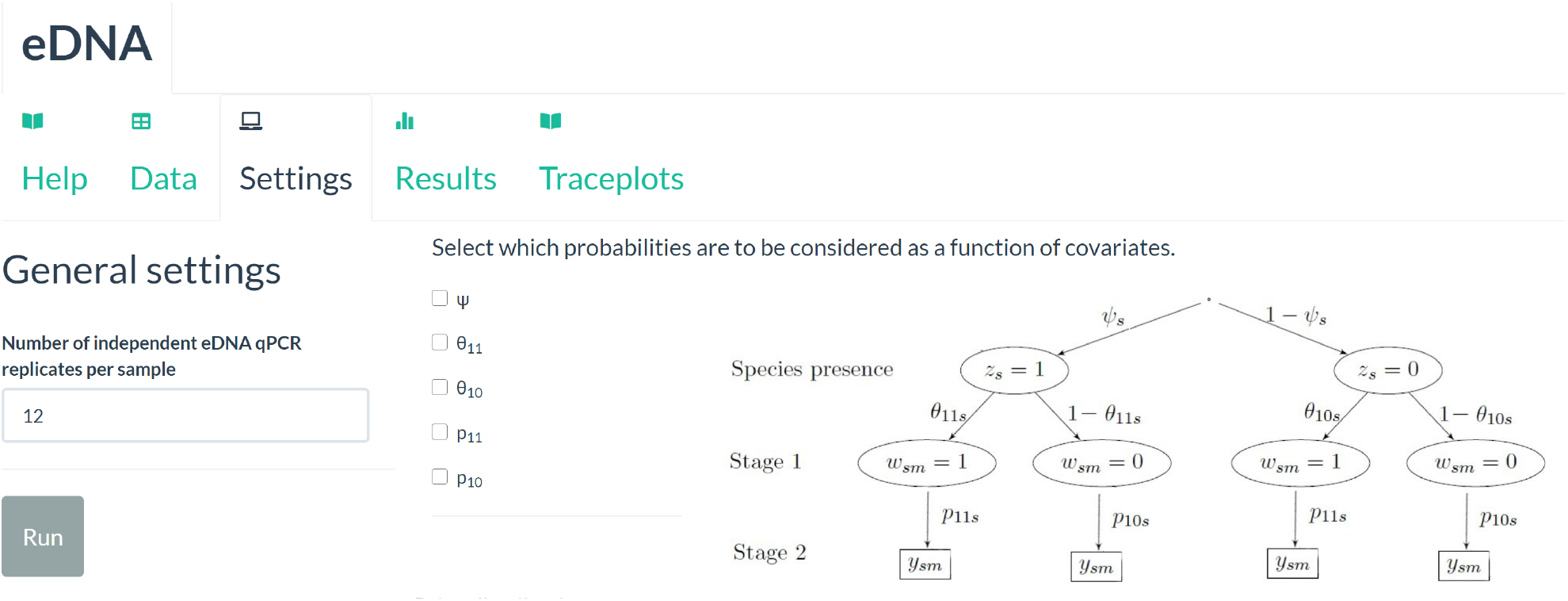
Settings tab in the eDNA RShiny app

The settings window also allows users to change the number of iterations, including number of chains, burn-in and thinning, as well as the prior distribution parameters, although we would recommend that the prior settings are not changed unless the user has a good understanding of the model. Once the user clicks on Run, the app will begin model fitting and the iteration number will be shown in the bottom right corner.

The results are available in the Results tab. These include posterior summaries for all model parameters, or corresponding coefficients of covariates for parameters that have been modelled as functions of covariates. All of the results and figures that are produced as part of the output can be downloaded.

The diagnostics tab produces traceplots for all model parameters as well as effective sample sizes (ESS) obtained. A message will appear to indicate if any of the parameters have ESS lower than 500 so a closer inspection of the traceplots and ESS outputted would help identify the parameters that are not mixing well.

## 4 Case study

We consider a data set on great crested newts collected by Natural England in 2018. *M* =1 water sample was collected from each of *S* = 2215 sites and *K* =12 qPCR runs were performed for each water sample. We have considered six categorical covariates and four continuous covariates (listed in Table 2) for the probability of occupancy, while all other parameters have been modelled as constant. We have also accounted for confirmed species presences that were available for 120 sites (see Figure 1 for description on how opportunistic data of this type are modelled). We run 1000 burn-in iterations and 2000 additional iterations with thinning set to 20. Fitting the model using the RShiny server took just under 2 hours, despite the large number of sites and considerable number of covariates considered. When run with fewer or no covariates the model fitting took only a few seconds to return results

**Table 2:** List of covariates considered as predictors for the probability of occupancy of great crested newts in the case study. These are based on the Habitat Suitability Index proposed by Oldham et al. (2000).

| Covariate | Type | Levels |
| --- | --- | --- |
| Geographic Location | Categorical | 1 - Optimal; 2 - Marginal; 3 - Unsuitable |
| Frequency of pond drying | Categorical | 1 - Never; 2 - Rarely; 3 - Sometimes; 4 - Annually |
| Water Quality | Categorical | 1 - Bad; 2 - Poor; 3 - Moderate; 4 - Good |
| Waterfowl intensity | Categorical | 1 - Absent; 2 - Minor; 3 - Major |
| Fish intensity | Categorical | 1 - Absent; 2 - Possible; 3 - Minor; 4 - Major |
| Terrestrial Habitat Quality | Categorical | 1 - Bad; 2 - Poor; 3 - Moderate; 4 - Good |
| Percentage Pond Shading | Numerical | NA |
| Pond Area | Numerical | NA |
| Pond Density | Numerical | NA |
| Percentage Macrophyte cover | Numerical | NA |

The app outputs posterior summaries of the site-specific probability of species presence, saved in a .csv file. For illustration purposes, we plot these summaries for a random sample of 50 sites in Figure 3 and provide the code for producing similar plots in the Help section of the app.

**Figure 3:**
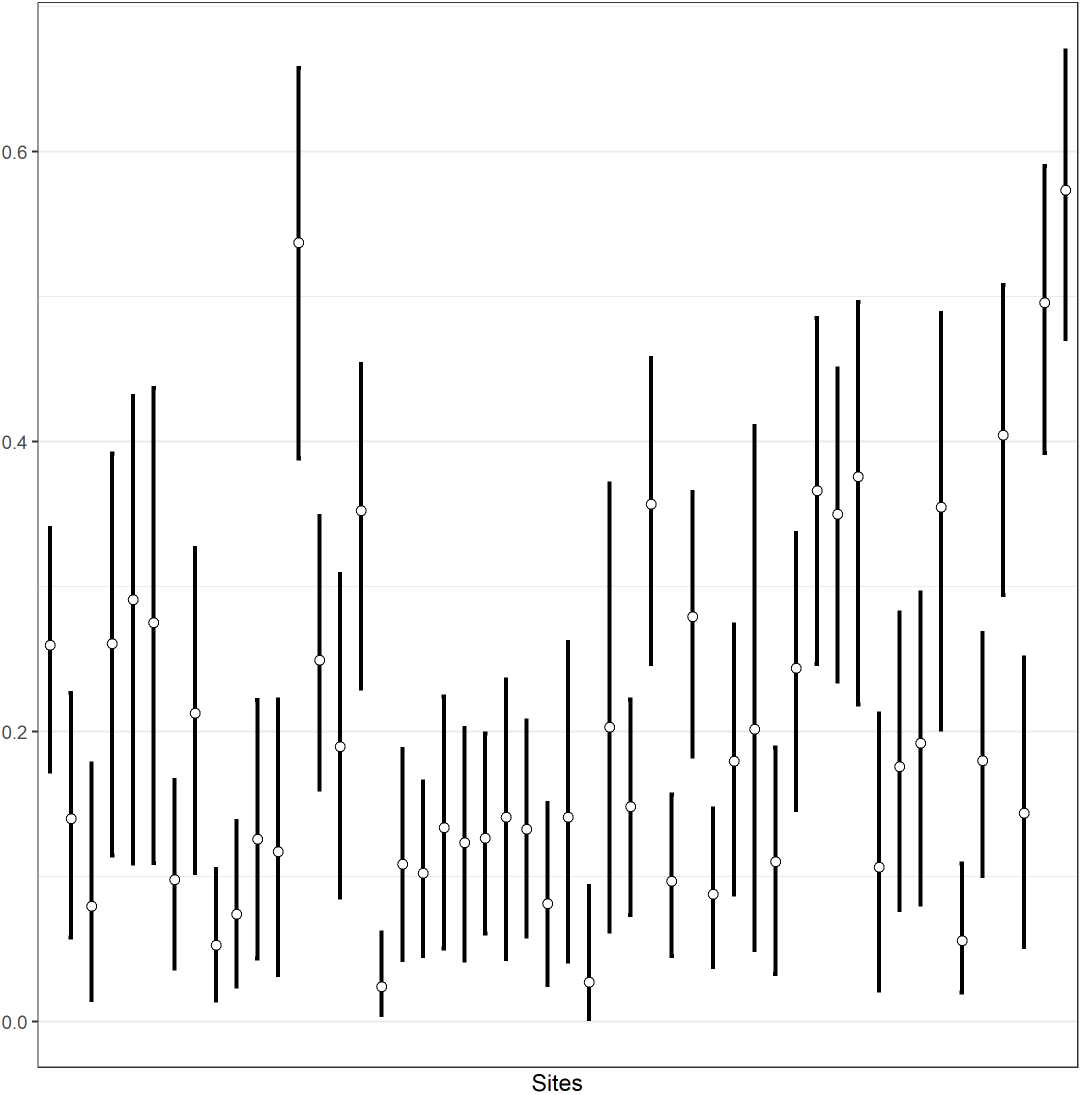
Posterior summaries of site-specific probabilities of occupancy for a random sample of 50 sites.

The PIPs for the probability of occupancy are provided by the app in a plot (see Figure 4). A high PIP indicates stronger support for a covariate as a predictor. In this case water quality, pond area, fish presence, percentage of macrophyte cover and frequency of pond drying all stand out as important predictors for the probability that a pond is occupied by great crested newts.

**Figure 4:**
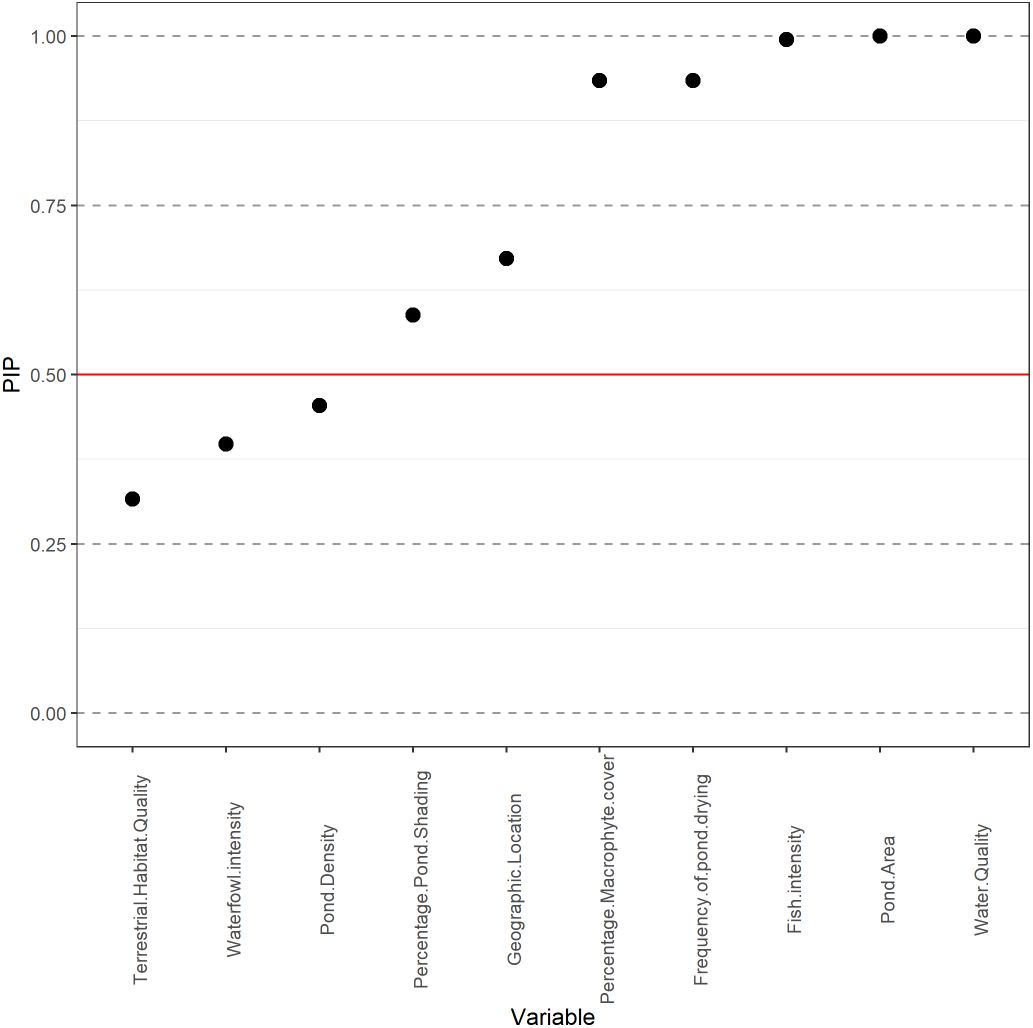
PIPs for the probability of occupancy. The horizontal line indicates the PIP=0.5 line.

Posterior summaries of the corresponding coefficients are given in Figure 5, showing that, even though geographic location and the percentage of pond that is shaded had PIPs above the 0.5 threshold, the 95% posterior credible intervals (PCIs) for the corresponding coefficients include 0. On the other hand, we can see that better water quality, lower pond area, lower levels of fish presence, higher levels of macrophytes and pond desiccation increase the probability of a pond being occupied by great crested newts. The predictors identified by the model and their corresponding effects are broadly consistent with current understanding of the preference of great crested newts for vegetated, fish-free and clean water ponds (Oldham et al., 2000). These results demonstrate that important predictors for the probability of species presence can be identified using eDNA data and our RShiny app.

**Figure 5:**
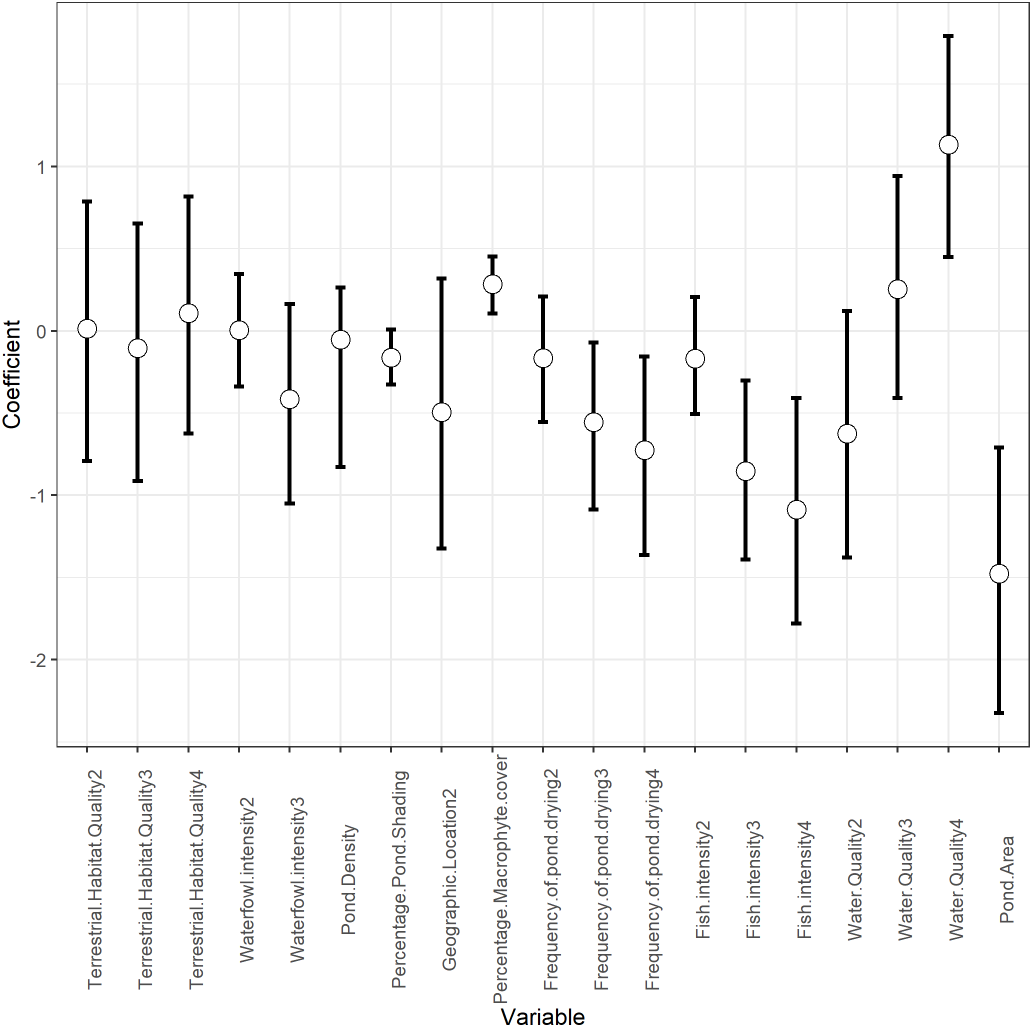
Posterior summaries of the coefficients of covariates for the probability of occupancy.

Posterior summaries of the probabilities related to observation error in both stages are given in Table 3. Stage 1 is related to higher probabilities of false observations, either positive or negative, compared to stage 2. The processes by which samples are collected mean that it is more likely for DNA to fail to be collected in the field, or contamination to be introduced at this stage, while lab protocols are more tightly controlled.

**Table 3:** Posterior summaries of the probabilities of a positive observation, true or false, in both stages of the survey.

| Parameter | Mean | 95% PCI |
| --- | --- | --- |
| $\theta_{11}$ | 0.867 | (0.802, 0.922) |
| $\theta_{10}$ | 0.078 | (0.004, 0.145) |
| $p_{11}$ | 0.862 | (0.851, 0.872) |
| $p_{10}$ | 0.026 | (0.023, 0.028) |

Finally, the app outputs the posterior probability of species absence conditional on *x* = 0,…,*K* positive qPCR replicates. For this example, the results are shown in the first row of Table 4 where we can see that the posterior conditional probability of species absence is very close to 1 given four or fewer qPCR positives, but then it declines sharply and plateaus at around 37%. The second row of Table 4 shows the posterior probability of *x, x* = 0,…, 12, positive qPCR replicates conditional on species presence. This conditional distribution is clearly bimodal. Specifically, the posterior probability of zero qPCR positives given species presence is just under 10% and this probability decreases for *x* = 1, 2, 3,4 before it starts to increase again reaching the second peak at *x* =11 (28%). This bimodality is due to the observation error in stage 1: the first peak at 0 is a result of a stage 1 false negative observation, whereas the second peak is a result of a stage 1 true positive observation.

**Table 4:** First row: Posterior probability of species absence conditional on *x* = 0,…, *K* positive qPCR replicates. Second row: Posterior probability of *x* positive qPCR replicated conditional on species presence.

|  | 0 | 1 | 2 | 3 | 4 | 5 | 6 | 7 | 8 | 9 | 10 | 11 | 12 |
| --- | --- | --- | --- | --- | --- | --- | --- | --- | --- | --- | --- | --- | --- |
| $1 - \psi(x)$ | 0.9766 | 0.9766 | 0.9766 | 0.9765 | 0.9462 | 0.4213 | 0.3683 | 0.3681 | 0.3681 | 0.3681 | 0.3681 | 0.3681 | 0.3681 |
| $q(x)$ | 0.0976 | 0.0308 | 0.0045 | 0.0004 | 0.0001 | 0.0003 | 0.0023 | 0.0123 | 0.0477 | 0.1319 | 0.2465 | 0.2798 | 0.1458 |

It is important to note that when *M* = 1 the model is non-identifiable and hence the results obtained are not reliable, unless the probability of occupancy, *ψ*, or the probabilities of observation error in stage 1, *θ*_11_ and *θ*_10_, are modelled as functions of covariates. Incorporating covariates helps overcome the identifiability issues, while an alternative solution is to incorporate information on confirmed species presences at some of the sites. Similarly, when *K* =1, the probabilities of observation error, either in stage 1 (*θ*_11_ and *θ*_10_) or in stage 2 (*p*_11_ and *p*_10_), need to be functions of covariates for the model to be identifiable.

## 5 Discussion

As eDNA surveys become increasingly used as monitoring tools, they have the potential to replace traditional survey methods that rely on direct observation of species, especially for difficult to detect species. Our RShiny app provides the necessary tool for researchers and practitioners to analyse their single-species eDNA data and obtain reliable estimates of site-specific probabilities of species presence while accounting for false positive and false negative observation error.

Unlike previous R-packages for fitting multi-scale occupancy models that have been applied to eDNA data (Dorazio and Erickson, 2018; Stratton et al., 2020), our implementation of the Griffin et al. (2019) model is novel in that it enables the estimation of false positive as well as false negative observation errors, both of which are known to be non-negligible in eDNA surveys. In addition, our RShiny app enables efficient Bayesian variable selection, which works well even when the number of predictors to be considered is large.

## Acknowledgements

We thank Natural England for collecting the data and making it available in an open access format. We also thank Diana Cole for useful insights about the issue of model identifiability. This work was partly funded by NERC, as part of project NE/T010045/1 “Integrating new statistical frameworks into eDNA survey and analysis at the landscape scale”.

## Authors’ contributions

AD developed the RShiny app, EM advised AD during the development of the app, tested the app and drafted the manuscript, JEG developed the original model and advised on implementing it within the app, ASB and RAG provided the data and interpretation of the results. All authors edited the manuscript and approved submission.

## Data availability statement

The example data set was collected by Natural England as part of a species distortion assessment project, and made available through the Natural England open data portal: https://naturalengland-defra.opendata.arcgis.com/datasets/ffba3805a4d9439c95351ef7f26ab33c_0/data.

## References

Brown, P. J., M. Vannucci, and T. Fearn (1998). Multivariate bayesian variable selection and prediction. Journal of the Royal Statistical Society: Series B (Statistical Methodology) 60(3), 627–641.

Chipman, H., E. I. George, R. E. McCulloch, M. Clyde, D. P. Foster, and R. A. Stine (2001). The practical implementation of bayesian model selection. Lecture Notes-Monograph Series, 65–134.

Dorazio, R. M. and R. A. Erickson (2018). ednaoccupancy: An r package for multiscale occupancy modelling of environmental dna data. Molecular ecology resources 18(2), 368–380.

Ghosh, J. (2015). Bayesian model selection using the median probability model. Wiley Interdisciplinary Reviews: Computational Statistics 7(3), 185–193.

Griffin, J. E., E. Matechou, A. S. Buxton, D. Bormpoudakis, and R. A. Griffiths (2019). Modelling environmental dna data; bayesian variable selection accounting for false positive and false negative errors. Journal of the Royal Statistical Society: Series C (Applied Statistics).

Guillera-Arroita, G., J. J. Lahoz-Monfort, A. R. van Rooyen, A. R. Weeks, and R. Tingley (2017). Dealing with false-positive and false-negative errors about species occurrence at multiple levels. Methods in Ecology and Evolution 8(9), 1081–1091.

McClenaghan, B., Z. G. Compson, and M. Hajibabaei (2020). Validating metabarcoding-based biodiversity assessments with multi-species occupancy models: A case study using coastal marine edna. PloS one 15(3), e0224119.

Oldham, R., J. Keeble, M. Swan, and M. Jeffcote (2000). Evaluating the suitability of habitat for the great crested newt (triturus cristatus). Herpetological Journal 10(4), 143–155.

Polson, N. G., J. G. Scott, and J. Windle (2013). Bayesian inference for logistic models using pólya–gamma latent variables. Journal of the American statistical Association 108(504), 1339–1349.

Stratton, C., A. J. Sepulveda, and A. Hoegh (2020). msocc: Fit and analyse computationally efficient multi-scale occupancy models in r. Methods in Ecology and Evolution.

Thomsen, P. F., J. Kielgast, L. L. Iversen, C. Wiuf, M. Rasmussen, M. T. P. Gilbert, L. Orlando, and E. Willerslev (2012). Monitoring endangered freshwater biodiversity using environmental dna. Molecular ecology 21(11), 2565–2573.

Thomsen, P. F. and E. Willerslev (2015). Environmental dna-an emerging tool in conservation for monitoring past and present biodiversity. Biological conservation 183, 4–18.

Valentini, A., P. Taberlet, C. Miaud, R. Civade, J. Herder, P. F. Thomsen, E. Bellemain, A. Besnard, E. Coissac, F. Boyer, et al. (2016). Next-generation monitoring of aquatic biodiversity using environmental dna metabarcoding. Molecular ecology 25(4), 929–942.

